# Evolution and mechanism of *MEIS2*-mediated forelimb specialization in bats

**DOI:** 10.64898/2026.06.03.729816

**Authors:** Bai-Wei Lo, Silvia Aldrovandi, David Meierhofer, Magdalena Schindler, Francisca Martinez Real, Alessa Ringel, Christian Feregrino, Stefan Mundlos

## Abstract

The genomic basis of limb adaptations in tetrapods is thought to be largely driven by changes in gene regulation. However, the mechanisms by which regulatory programs evolve are not well understood. In bats, wing membrane development has been shown to be associated with expression of the transcription factor *MEIS2* in the interdigital tissue of the forelimb. However, *MEIS2* alone is insufficient to recapitulate wing morphology, suggesting that its regulatory context has also undergone divergence. Here, we integrate functional genomics with sequence-to-function deep learning to dissect both the mechanistic and evolutionary roles of *MEIS2* in bat forelimb development. Using models trained on embryonic limb data from bat and mouse, we identify a strong association between *MEIS2* binding and the transcription factor *TWIST1*, a finding which is supported by single-cell transcriptomic analyses. To investigate the evolutionary dimension, we applied these models across more than 100 genomes, including extant bats, closely related species, and reconstructed ancestors. This analysis identified divergences in regulatory regions, which likely contribute to bat-specific forelimb expression of genes that lead to wing morphogenesis. Notably, these changes are prominent in the regulatory domains of the MEIS dimerization partner *PBX1*, indicating coordinated regulatory evolution. Together, our results demonstrate that the evolution of a complex morphological trait involves coordinated changes in both trans-regulatory environments and cis-regulatory landscapes. More broadly, this study provides a framework for integrating deep learning with comparative and functional genomics to investigate regulatory evolution.

## Introduction

Understanding how complex traits arise during macroevolution is a long-standing central question in biology. The morphological diversification of tetrapod limbs provides numerous parallel opportunities to investigate this question. Molecularly, tetrapod limb development follows a deeply conserved blueprint involving well-known signaling pathways and transcription factors (Petit et al. 2017). Studies on tetrapod lineages with diverse limb morphology have revealed that important contributions from alterations in gene regulation during development play an important role in modifying the ancestral blueprint (Cooper et al. 2014; Lopez-Rios et al. 2014; Eckalbar et al. 2016; Kvon et al. 2016; Leal and Cohn 2018; Young et al. 2019). However, the precise mechanisms underlying these regulatory changes, and their broader impact on the evolution of associated regulatory regions, remain largely unexplored.

The evolution of bats (order Chiroptera), the most recently evolved group of tetrapods to acquire powered flight, is one of the most striking examples of morphological innovation in mammals. Bat forelimbs are drastically enlarged and transformed into wings, consisting of a thin membrane supported by elongated digits. Previous studies on the molecular basis of bat wing development have revealed the distinctive expression pattern of the transcription factor (TF) *MEIS2* during bat forelimb development (Dai et al. 2014; Mason et al. 2015; Lyu et al. 2025; Schindler et al. 2025). The homeobox TF *MEIS2*, is expressed in the proximal part of the early limb bud (Delgado et al. 2021), but becomes highly expressed in the later interdigital tissue of developing bat forelimbs, which eventually gives rise to the interdigital wing membranes (Figure 1) (Dai et al. 2014; Mason et al. 2015; Lyu et al. 2025; Schindler et al. 2025). Analyses of the functional genome and regulatory networks suggest that *MEIS2* activates multiple developmental genes specifically in the developing interdigital tissue, positioning it as a key regulator of wing membrane development (Schindler et al. 2025). However, transgenic assays have shown that the ectopic expression of *MEIS2* is not sufficient to fully reproduce the bat wing phenotype in a mouse model, despite recapitulating part of its transcriptional program (Schindler et al. 2025). This indicates divergence of additional aspects in *MEIS2* operation between species, which might involve modifications in *cis*-regulatory elements and *trans*-regulatory environments. Therefore, the details of the regulatory activity and mechanism of *MEIS2* have to be investigated further.

**Fig. 1.**
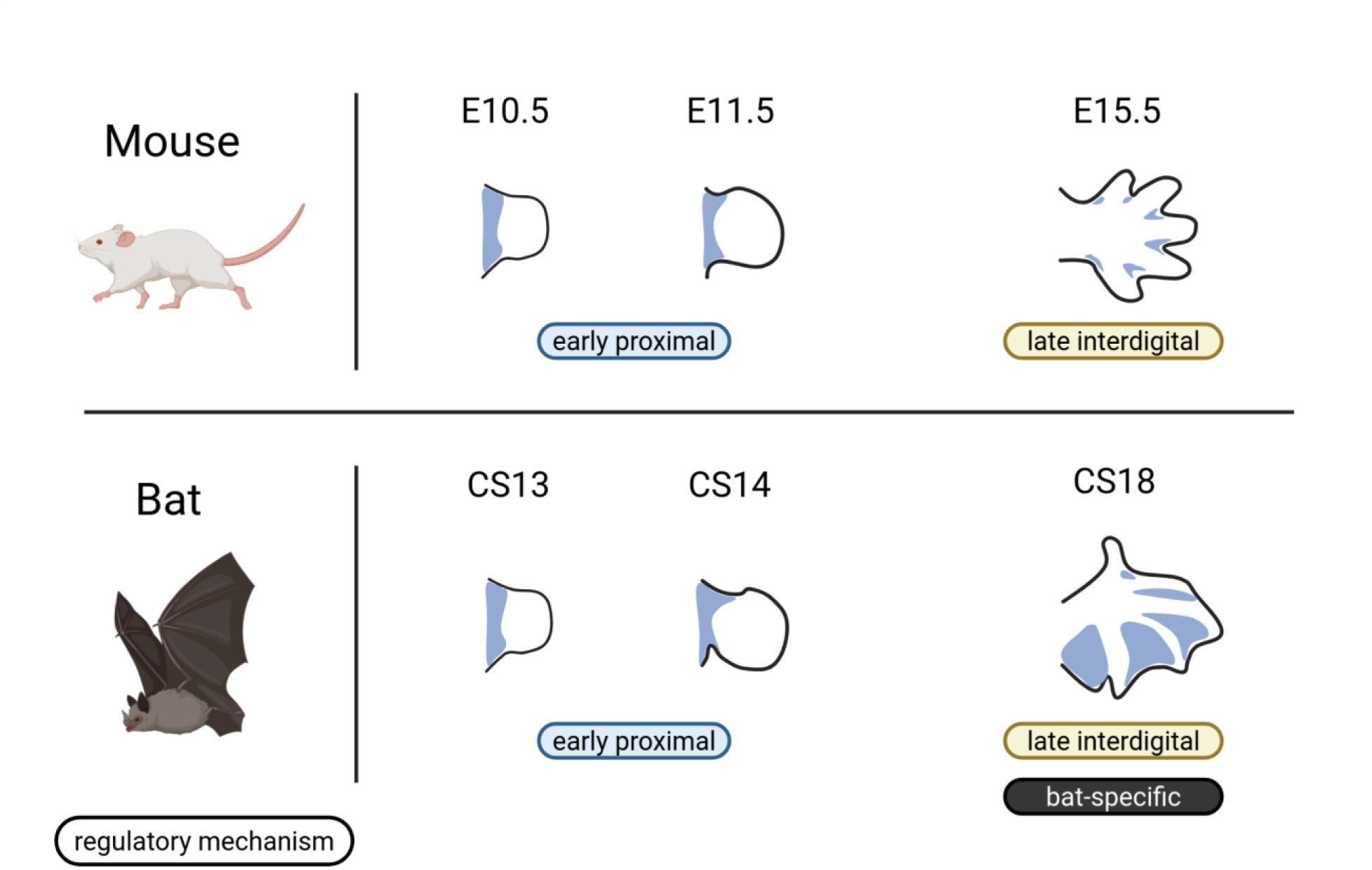
Forelimb MEIS expression patterns in mouse and bat and the research framework used to compare the mechanism underlying their differences. *MEIS1* and *2* are expressed proximally during early developmental stages. In later stages when bat forelimb becomes morphologically distinctive, *MEIS2* is expressed in the distal interdigital regions. The distinctive expression pattern in bat forelimb is likely a collective effect of a conserved late interdigital regulatory mechanism, which seem to be present also in mice, and additional novel bat-specific mechanisms. E11.5, E15.5, CS14, and CS18 *MEIS2* expression patterns are adapted from (Dai et al. 2014); E10.5 and CS13 (presumed) are adapted from (Delgado et al. 2021).

Recent advances in the field of deep learning algorithms provide powerful tools to aid in deciphering the regulatory genome (Zhou and Troyanskaya 2015; Kelley et al. 2016; Avsec et al. 2021; de Almeida et al. 2022; Vaishnav et al. 2022). Although these approaches are widely used to study human genetic variation, they can also be applied to help resolve macroevolutionary questions (Kaplow et al. 2023; Sarropoulos et al. 2026). Here, we integrate multi-omic analyses with deep learning-based predictions to systematically study the regulatory biology of the TF *MEIS2* during wing membrane development. We follow the classical “level of analysis” framework (Sherman 1988; Mayr 1993), addressing both proximate mechanistic and ultimate evolutionary dimensions. At the proximate level, we aimed to identify bat-specific regulatory mechanisms linked to *MEIS2* DNA-binding. Through deep learning analyses of epigenomic data, supported by single cell transcriptomic observations, we find that *MEIS2* DNA-binding is associated with the transcription factor *TWIST1*. At the ultimate level, we explored genome-wide evolutionary changes associated with *MEIS2* DNA-binding. We discovered that transposable elements contribute to novel regulatory elements in bat. Furthermore, by using our trained deep learning models to evaluate genomes from bats and related species, as well as reconstructed ancestral genomes, we infer bat-specific functional genomic changes in *MEIS2*-related regulatory elements.

Together, our findings provide new insights into the intricate molecular processes surrounding regulatory evolution, involving coordination of both *trans*-environments and *cis*-regulatory landscapes. This offers a generalizable framework for applying deep learning strategies to understand regulatory genomic changes in evolution.

## Results

### A cross species, cross stage study of *MEIS2* regulatory mechanisms in the bat forelimb

To better understand the evolutionary molecular basis of the *MEIS2*-mediated specialization of bat forelimbs, we compared the regulatory activity of *MEIS2* across different species. To characterize the *MEIS2* DNA-binding landscape in the interdigital tissues of developing bat forelimbs, we conducted MEIS ChIP-seq analysis on embryonic autopod forelimbs of Seba’s short-tailed bat (*Carollia perspicillata*) at stage CS18 (Figure 1) (Cretekos et al. 2005). For comparison, we performed MEIS ChIP-seq analysis on the autopod limbs of mice at the corresponding stage (E15.5, Fig. 1). This revealed that the bat genome exhibits approximately 9,800 significant MEIS binding peaks in this tissue, whereas in mice, detectable binding peaks at this stage are scarce in both fore- and hindlimb (< 1000). This aligns with the observed differences in *MEIS2* expression levels between the two species. However, the limited number of peaks in mice prevents the use of common statistical approaches (e.g. motif enrichment) for cross-species comparisons at this developmental stage.

In contrast to the bat’s distinct interdigital expression of *MEIS2*, its limb bud expression in earlier developmental stages resembles the proximal pattern observed in mice and other tetrapods (Fig. 1) (Mercader et al. 1999; Mercader et al. 2005; Dai et al. 2014). Comparing conserved early proximal and divergent late interdigital *MEIS2* DNA-binding landscapes is an opportunity to clarify stage-specific regulatory mechanisms, and potentially reveal evolutionary adaptations specific to the bat forelimb (Fig. 1). Obtaining early-stage samples from *C. perspicillata* is challenging, making direct temporal comparisons along bat development difficult. However, since the early proximal pattern is conserved in both species, the mouse binding landscape might represent a shared mammalian archetype. Thus, we used a published early E10.5 mouse limb ChIP-seq dataset (Delgado et al. 2021) to perform a cross-stage comparison and investigate developmental and evolutionary changes in bats (early, proximal, murine, versus late, interdigital, chiropteran). Reanalyzing the available data, we identified 3,200 MEIS binding peaks in the early E10.5 mouse forelimb (referred to as MEIS binding peaks since *MEIS1* and *2* paralogs can not be distinguished). This interspecific and interstage epigenomic dataset provides a framework for investigating evolutionary shifts in MEIS binding associated with bat forelimb development.

### Conserved MEIS DNA-binding properties in the developing limb across species

We next asked whether MEIS homologs share similar DNA-binding properties in bat and mouse limbs. Motif enrichment analysis of MEIS peaks from bat late-interdigital tissue and mouse early-proximal limb tissue revealed enrichment for highly similar motifs in both species, with MEIS and PBX motifs being the most significantly enriched (Fig. 2A, Supplementary Tables 1–4). However, direct comparison of motif enrichment results across developmental stages or species can be challenging as the analysis provides only summary-level statistics. To directly assess MEIS DNA-binding potential, we therefore turned to sequence-to-function machine learning methods. We trained a binary deep learning classifier using bat MEIS ChIP-seq peaks and ATAC-seq–defined accessible chromatin (Fig. 2B). This bat MEIS binding model accurately distinguished MEIS-bound from unbound accessible regions in the *C. perspicillata* late forelimb (ROC AUC = 0.91; Fig. 2C).

**Fig. 2.**
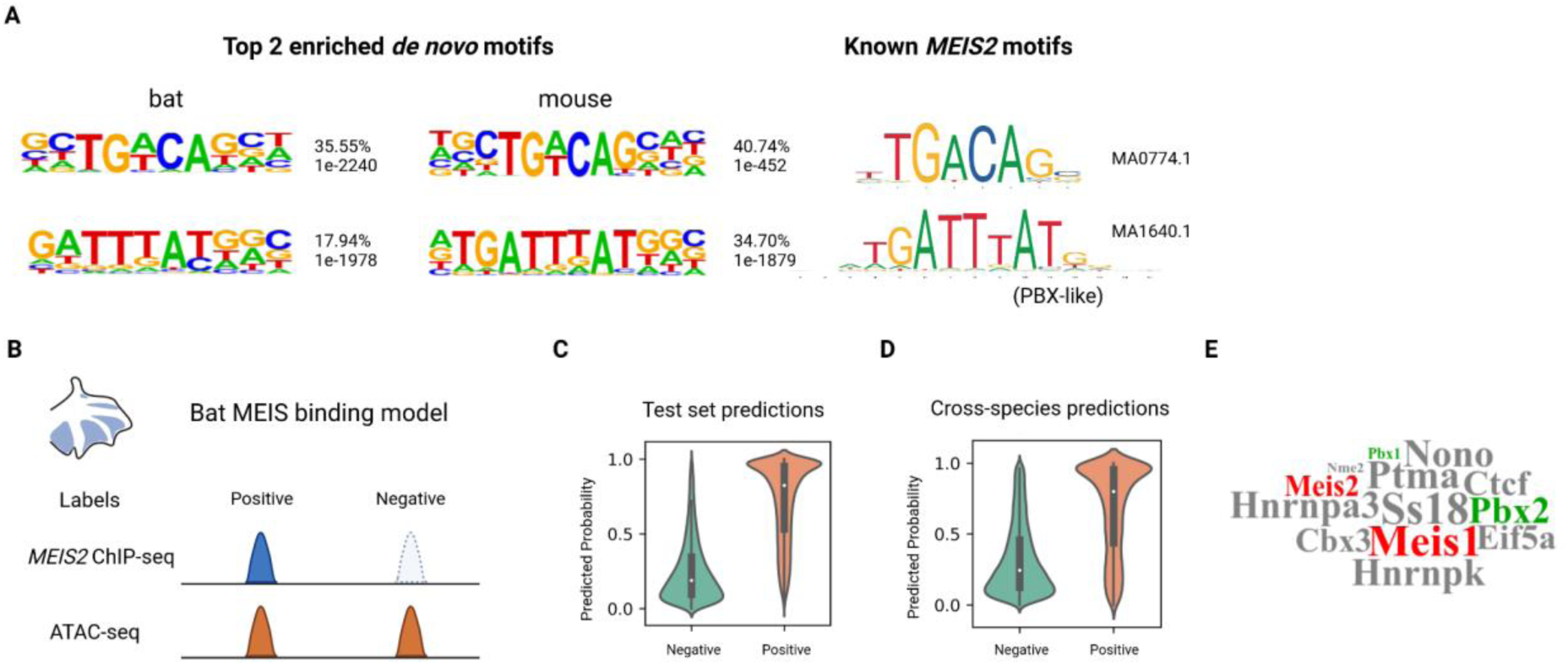
Shared MEIS binding properties in bat and mouse. **A.** *De novo* enriched motifs from bat and mouse MEIS ChIP-seq. Percentage of peaks containing the motif and q values are shown next to the motifs. MA1640.1 has been proposed to be bound by PBXs, which forms a heterodimer with MEIS factors (Penkov et al. 2013). **B.** Design of bat MEIS binding model. ATAC-seq from CS17 bat forelimb was used to identify accessible regions. **C.** Prediction of bat MEIS binding model on held out test chromosomes. **D.** Prediction of bat MEIS binding model on mouse limb MEIS peaks and non-peak accessible regions. **E.** Proteins that co-purified with *MEIS2* in bat CS18 forelimb. The sizes of the words are based on log sample/control values. Besides PBXs, no other activating transcription factors were detected.

To test whether MEIS exhibits similar DNA-binding properties in the mouse limb, we analyzed ATAC-seq data from E10.5 mouse forelimbs and evaluated these regions using the bat-trained model. The model successfully predicted MEIS-bound peaks within accessible mouse limb chromatin (ROC AUC = 0.81; Fig. 2D), suggesting similar MEIS-binding properties across stages and species.

In addition to functional genomic analyses, we searched for MEIS-interacting proteins in the bat late-interdigital tissue using rapid immunoprecipitation–mass spectrometry of endogenous proteins (RIME). RIME identified MEIS itself and several chromatin modifiers such as *EIF5A* or *SS18* (Fig. 2E). *PBX1* and *2* were identified as the only relevant interacting TFs, reinforcing the canonical role of PBXs as main co-factors in MEIS function also in the interdigital forelimb (Chang et al. 1997; Shanmugam et al. 1999; Delgado et al. 2021). Together, our machine learning and proteomic results indicate that key MEIS DNA-binding properties, including binding motifs and the MEIS co-factor PBX, are conserved across species and developmental stages within limb mesenchymal tissues. Interestingly, besides PBXs, RIME did not detect any other activating TFs that form complexes with *MEIS2* (Fig. 2E). This suggests more subtle interactions between *MEIS2* and other functionally relevant TFs in the bat interdigital forelimb.

### *TWIST1* or similar bHLH factors are involved in interdigital MEIS binding in the bat

We next compared the forelimb MEIS DNA-binding landscapes between early-proximal mouse limbs and late-interdigital bat limbs. To ensure reliable homology inference, we restricted our analysis to regions that can be aligned between genomes (Fig. 3A). These regions are remarkably conserved, even across vertebrates (Supplementary Fig. 1). Strikingly, despite this high level of sequence conservation, most MEIS-bound peaks within these regions are species/stage-specific (91% in bat and 88% in mouse, Fig. 3A).

**Fig. 3.**
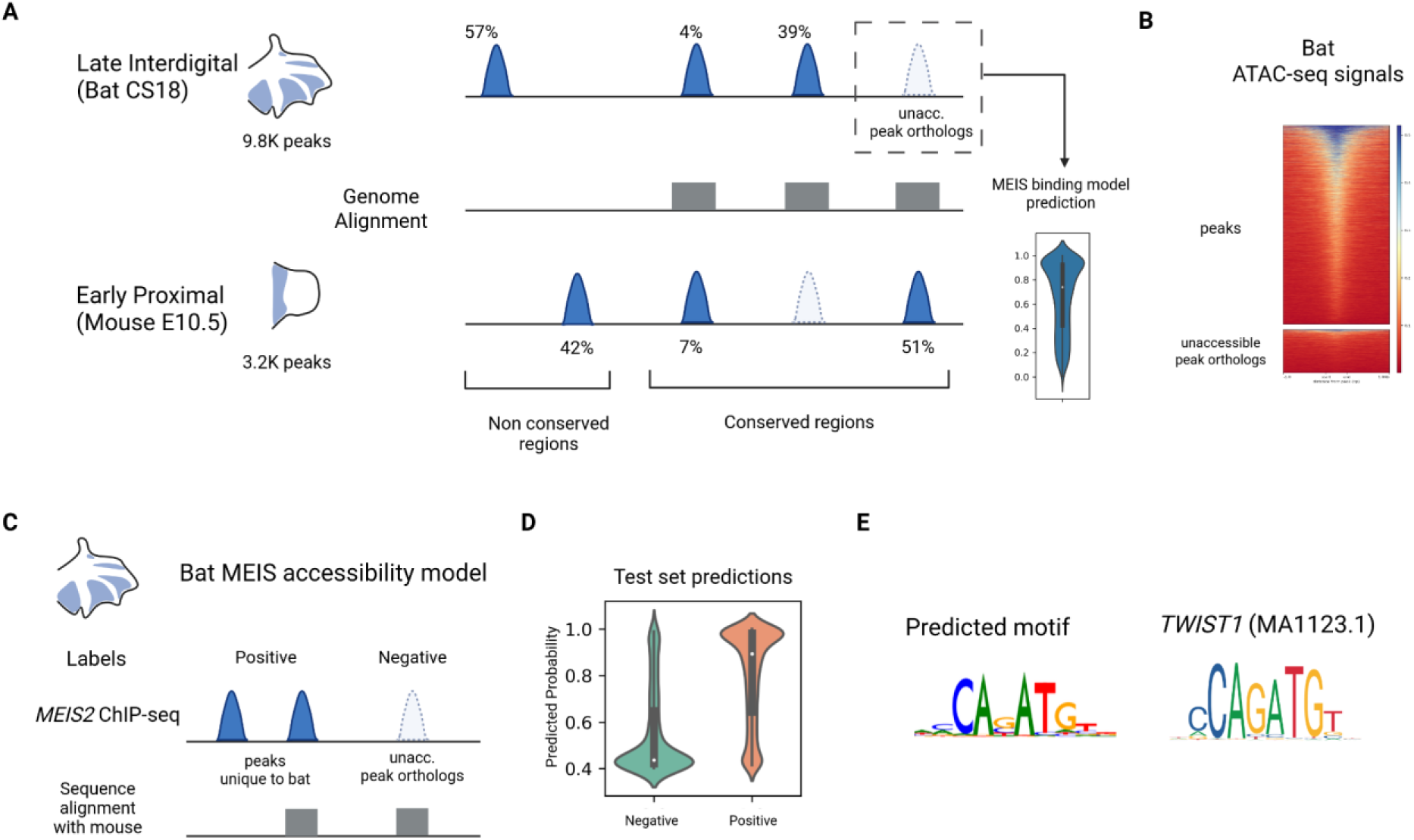
Late interdigital MEIS accessibility in bat is associated with a *TWIST1*-like motif. **A.** Cross-stage comparison of MEIS binding landscape between bat and mouse. The peak orthologs from the carPer2 genome showed overall high predicted MEIS binding affinities. **B.** ATAC-seq accessibility signals in bat MEIS-binding peaks and inaccessible peak orthologs. The center 501 bp region or a peak summit, and the 1k bp flanking regions, are shown in the figure. **C.** Design of accessibility model. The binary classification model differentiates late interdigital bat-specific peaks from unacc. peak orthologs. **D.** Prediction values of accessibility model on held out test chromosomes. **E.** Model interpretation by TF-modisco discovered a motif that highly resembles the canonical motif of *TWIST1* (MA1123.1).

We hypothesized that both the bat-specific peaks and the non-binding mouse peak orthologs in bat represent *bona fide* MEIS binding elements, but that their differential occupancy is driven by differences in chromatin accessibility, since accessibility is a prerequisite for transcription factor binding. Consistent with this hypothesis, bat MEIS peaks exhibit significantly higher (Mann-Whittney U test p < 0.01) ATAC-seq signals than the non-binding orthologs of mouse peaks (“unacc. peak orthologs”; Fig. 3B).

To further evaluate this hypothesis, we used the bat MEIS binding model—which predicts binding affinity in accessible regions—to assess the binding potential of unacc. peak orthologs. These regions displayed binding score distributions similar to *bona fide* MEIS-bound peaks (Fig. 3A, Fig. 2D-E), indicating that they retain strong intrinsic MEIS binding potential. Together, these results support a model in which differential chromatin accessibility governs MEIS occupancy within conserved genomic regions in different developmental time and space.

To investigate the mechanisms associated with differential accessibility, we trained a second deep learning model: the bat MEIS accessibility model (Fig. 3C). This model distinguishes MEIS-bound peaks from unacc. peak orthologs in the bat genome (ROC AUC = 0.81; Fig. 3D), effectively predicting late-interdigital-specific, or “wing-specific,” accessibility. Using TF-modisco to interpret the model and identify motifs driving these accessibility predictions, we found that a *TWIST1*-like motif had the highest attribution for predicting wing-specific accessibility of MEIS-binding regions (Fig. 3E).

### Overlapping binding of *MEIS2* and *TWIST1* in bat interdigital tissues

We next investigated the interactions between MEIS and *TWIST1* in limb mesenchymal tissues. By reanalyzing a published dataset (Kim et al. 2024), we discovered that 18.91% of *MEIS* binding peak summits are located within 50 base pairs of a *TWIST1* ChIP-seq peak summit in E10.5 mouse forelimbs, indicating proximal binding between these transcription factors (Fig. 4A). Given the emphasis of the MEIS accessibility model on *TWIST1’s* importance for wing-specific MEIS binding, we anticipate an even higher density of DNA-bound *TWIST1* within MEIS peaks in developing bat interdigital tissues. We reexamined the enriched motifs and found significant enrichment of both canonical (Fig. 3E) and a *TWIST1*-homeodomain composite motifs in bats (Supplementary Table 3) (Kim et al. 2024). To further study *TWIST1* DNA-binding within individual MEIS peaks, and in the absence of bat *TWIST1* ChIP-seq data, we developed a Mouse *TWIST1* binding model using E10.5 mouse limb data (Fig. 4B). This model is conceptually similar to the bat MEIS binding model and successfully predicts *TWIST1* DNA-binding in mouse limb accessible genomic regions with a ROC AUC of 0.92 (Fig. 4C). The model also recapitulated the within-50-bp proximal binding between *TWIST1* and MEIS (ROC AUC = 0.79). As a control, analysis of MEIS ChIP-seq peaks from adult mouse heart tissue, which does not express *TWIST1*, resulted in low *TWIST1* DNA-binding predictions (Fig. 4D).

**Fig. 4.**
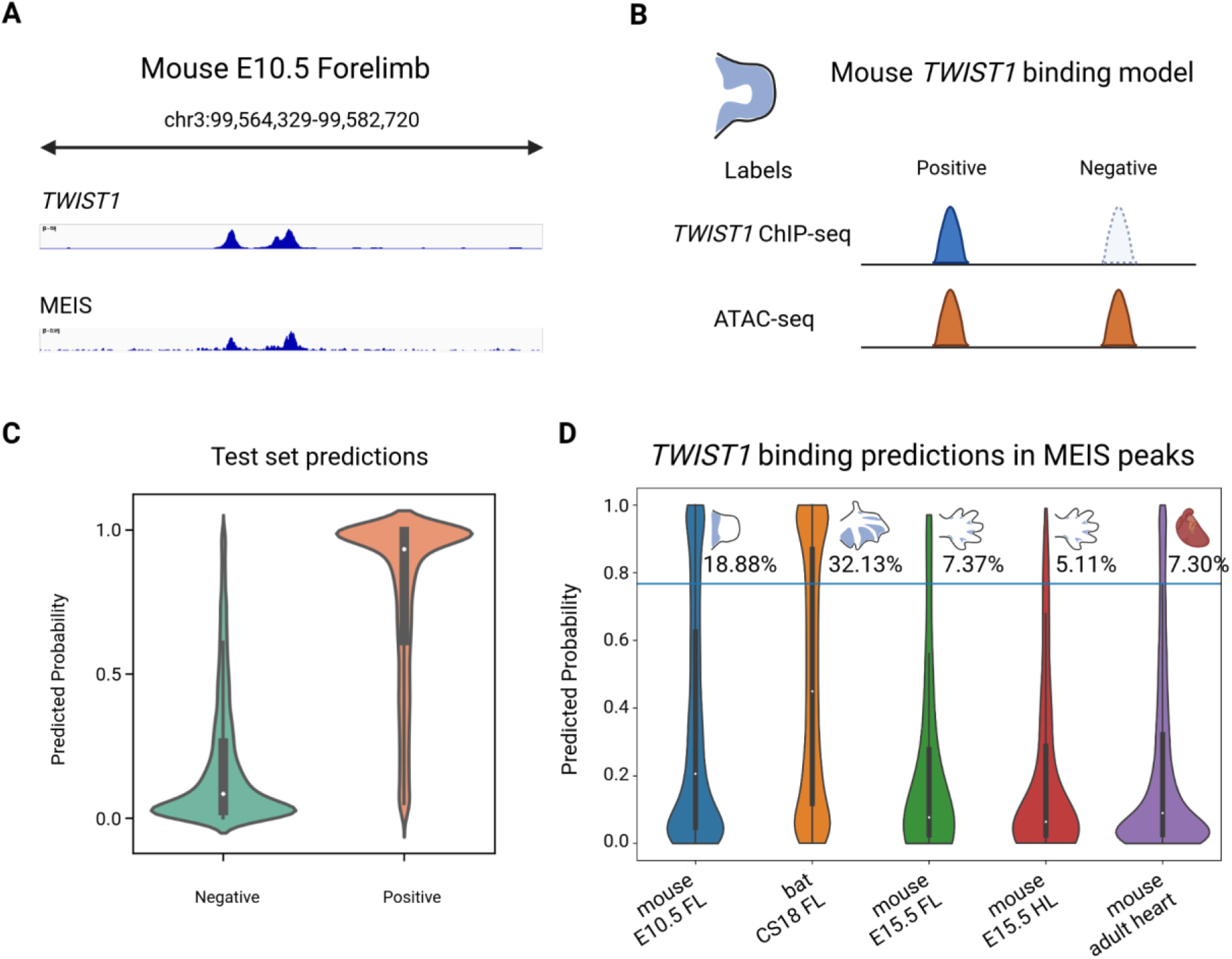
A Mouse *TWIST1* binding model reveals that many *MEIS2* binding sites are shared by *TWIST1* in the bat interdigital tissue. **A.** ChIP-seq tracks showing examples of close binding between *TWIST1* and MEIS in E10.5 mouse forelimb. **B.** Design of Mouse *TWIST1* binding model. *TWIST1* expression pattern is adapted from (Firulli et al. 2005). **C.** Prediction of *TWIST1* binding model on held out test chromosomes. **D.** Prediction of *TWIST1* binding in the MEIS binding peaks of: mouse E10.5 forelimb, bat CS18 forelimb, mouse E15.5 forelimb, mouse E15.5 hindlimb, mouse adult heart. The numbers represent the percentage of peaks with prediction values larger than the blue line (0.767). This cutoff value was chosen based on the ground truth of *TWIST1* and MEIS ChIP-seq overlapping in the E10.5 mouse forelimb. FL: forelimb; HL: hindlimb.

We then applied the *TWIST1* binding model to MEIS peaks from various limb tissues in bat and mouse: CS18 bat forelimb, E15.5 mouse fore-, and hindlimb (Fig. 1). In line with the previous predictions from the MEIS accessibility model, 32.13% of late interdigital bat peaks were predicted to be bound by *TWIST1* (Fig. 4D), significantly higher than early proximal mouse peaks (18.88%, p < 0.05). Furthermore, predicted *TWIST1* binding in late interdigital MEIS peaks in both mouse forelimb and hindlimb are low and comparable to that in the adult heart (Fig. 4D). However, it is important to interpret these results with caution as they are based on predictions from fewer than 1,000 peaks in late mouse stages. In summary, the high predicted proportion of *TWIST1* binding in bat forelimb MEIS peaks compared to mouse underscores *TWIST1’s* role in bat-specific regulatory mechanisms for interdigital MEIS binding (Fig. 1).

### Single-cell transcriptome verifies bat-specific upregulation of *TWIST1* in *MEIS2*+ cells in the forelimb

Based on our epigenomic and deep learning analyses, we anticipated bat-specific expression patterns of *TWIST1* in the developing forelimbs. To verify this, we analyzed a single cell transcriptomic atlas of embryonic limbs from mouse and bat (Schindler et al. 2025). For each species, we extracted distal fore- and hindlimb fibroblasts expressing *MEIS2*, which are most likely of interdigital origin (Fig. 1). We found that *TWIST1* is indeed expressed in bat forelimb *MEIS2*+ cells (Fig. 5A).

**Fig. 5.**
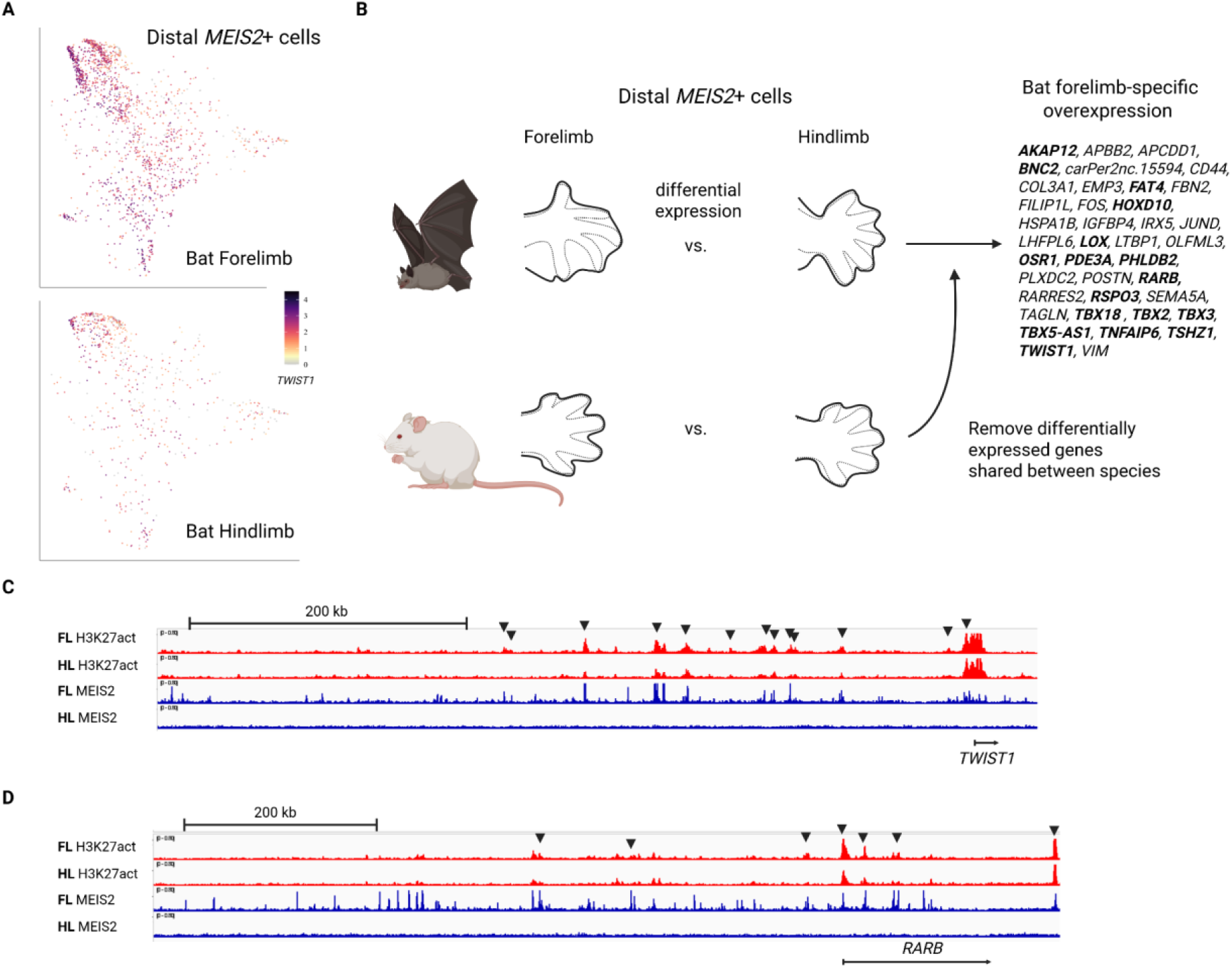
Single cell expression analyses of distal *MEIS2*+ cells and regulatory signals in differentially expressed genes. **A.** UMAPs showing expression levels of *TWIST1* in distal *MEIS2*+ cells in bat limbs. **B.** Identifying bat forelimb-specific overexpressed genes from a cross species single cell transcriptome atlas. Genes that have been reported in previous bat forelimb studies are highlighted in bold (Wang et al. 2010; Dai et al. 2014; Wang et al. 2014; Eckalbar et al. 2016; Maier et al. 2017; Schindler et al. 2025). **C.** and **D.** Differential limb regulatory signals in the *TWIST1* (**C.**) and *RARB* (**D.**) regulatory domain of bats. Bulk ChIP-seq from bat forelimb and hindlimb autopods reveal forelimb-differential H3K27act signals (black arrow heads) coupled with drastic difference in *MEIS2* binding signals. FL: forelimb; HL: hindlimb

To identify bat-specific expressional changes in the forelimb distal *MEIS2+* cells, we performed differential expression analysis between fore- and hindlimb cells within each species (log2FC > 0.25, adj. p value < 0.1). We then removed the genes that are differentially expressed between fore- and hindlimbs in both bat and mouse, leaving only 39 genes that are distinctively upregulated in the forelimb of bats (Fig. 5B). Notably, *TWIST1* and *RARB* are among these genes (Fig. 5A & B). The detection of *RARB*, a known marker of developing wing membranes (Mason et al. 2015), confirms that our approach indeed captures signals from the interdigital region.

To identify regulatory regions that may drive the differential expression of these genes, we analyzed bulk H3K27act (an epigenetic mark of active chromatin) ChIP-seq from the autopods of bat CS19 forelimbs and hindlimbs (Schindler et al. 2025). Within the topologically associating domains (TADs) of these 39 forelimb upregulated genes, we discovered 337 forelimb differentially active regions, over 50% of them overlap with forelimb *MEIS2* binding peaks (e.g. Fig. 5C & D). We performed gene ontology (GO) term enrichment analysis on the 39 genes, which revealed enriched terms related to transcription factor binding, tissue development, and extracellular matrix formation (Supplementary Table 5). Our scRNA-seq analysis confirms *TWIST1*’s co-expression with *MEIS2* in interdigital tissue. Moreover, it provides evidence of bat-specific *trans* evolution in the forelimb through upregulation of *TWIST1* and other TFs, while offering candidate *cis*-regulatory regions underlying such changes. The results, aligned with prior analyses, support *TWIST1*’s involvement in *MEIS2*-mediated interdigital wing membrane development in bats.

### Transposable elements contribute to *MEIS2* DNA-binding in bat forelimb development

We found that 57% of late interdigital *MEIS2* binding peaks in bats occur in genomic regions that are not conserved relative to mouse (Fig. 3A). Because non-conserved regions are often enriched for transposable elements (TEs), we examined TE content across conserved and non-conserved *MEIS2* peaks. Annotation of the *C. perspicillata* genome revealed that TEs overlap 38.6% of non-conserved *MEIS2* peaks, compared with only 9.6% of conserved peaks (χ² test, p < 0.001; Fig. 6A). To assess the functional contribution of TEs to MEIS binding, we disrupted TE sequences within non-conserved peaks by randomly shuffling their nucleotides and then predicted MEIS binding affinity using the MEIS binding model. Following TE disruption, 71.16% of peaks exhibited decrease in predicted MEIS binding, whereas only 27.21% showed an increase (Fig. 6B). Finally, we examined the TE class composition of peaks showing reduced binding after disruption, but found that no particular class stood out compared to the genome-wide TE distribution (Supplementary Fig. 2). These results indicate that transposable elements broadly and functionally contribute to *MEIS2* binding in bat forelimb development.

**Fig. 6.**
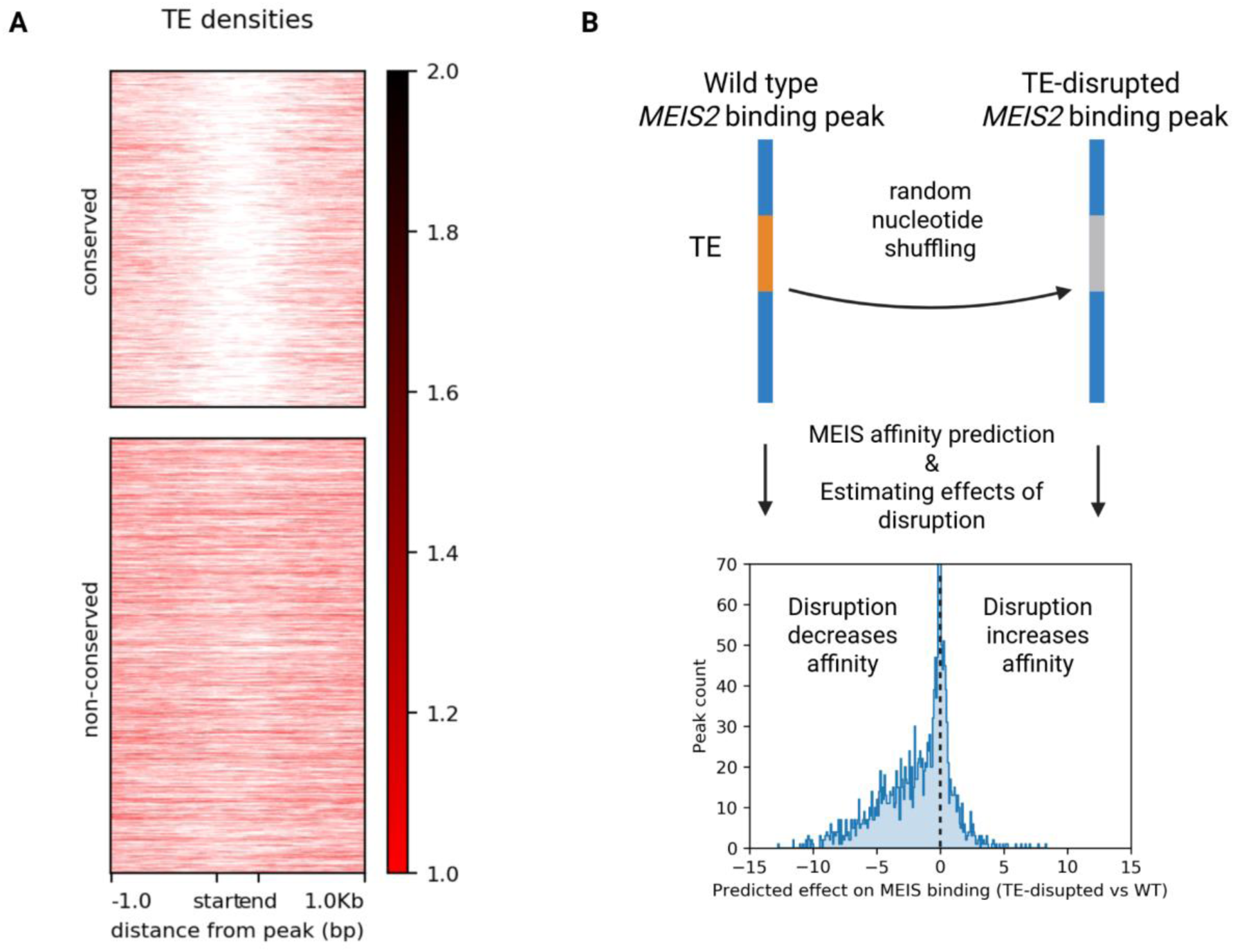
Abundance and impact of transposable elements on MEIS DNA-binding. **A.** *MEIS2*-binding peaks within conserved genomic regions are deprived of TEs. **B.** In general, disruption of TE sequences in peaks within non-conserved regions lowers the predicted MEIS binding affinity.

### Chiroptera-specific accessibility divergences in loci regulating *MEIS2*-interacting genes

Our machine learning models enable direct prediction of functional molecular phenotypes, such as wing-specific chromatin accessibility and transcription factor binding affinity, from DNA sequence, allowing functional comparisons across species without experimental profiling. By predicting these phenotypes at orthologous loci in genomes of bats and close relatives from the Zoonomia database (Armstrong et al. 2020), we aim at identifying regulatory regions with bat-specific divergence patterns (Fig. 7A).

**Fig. 7.**
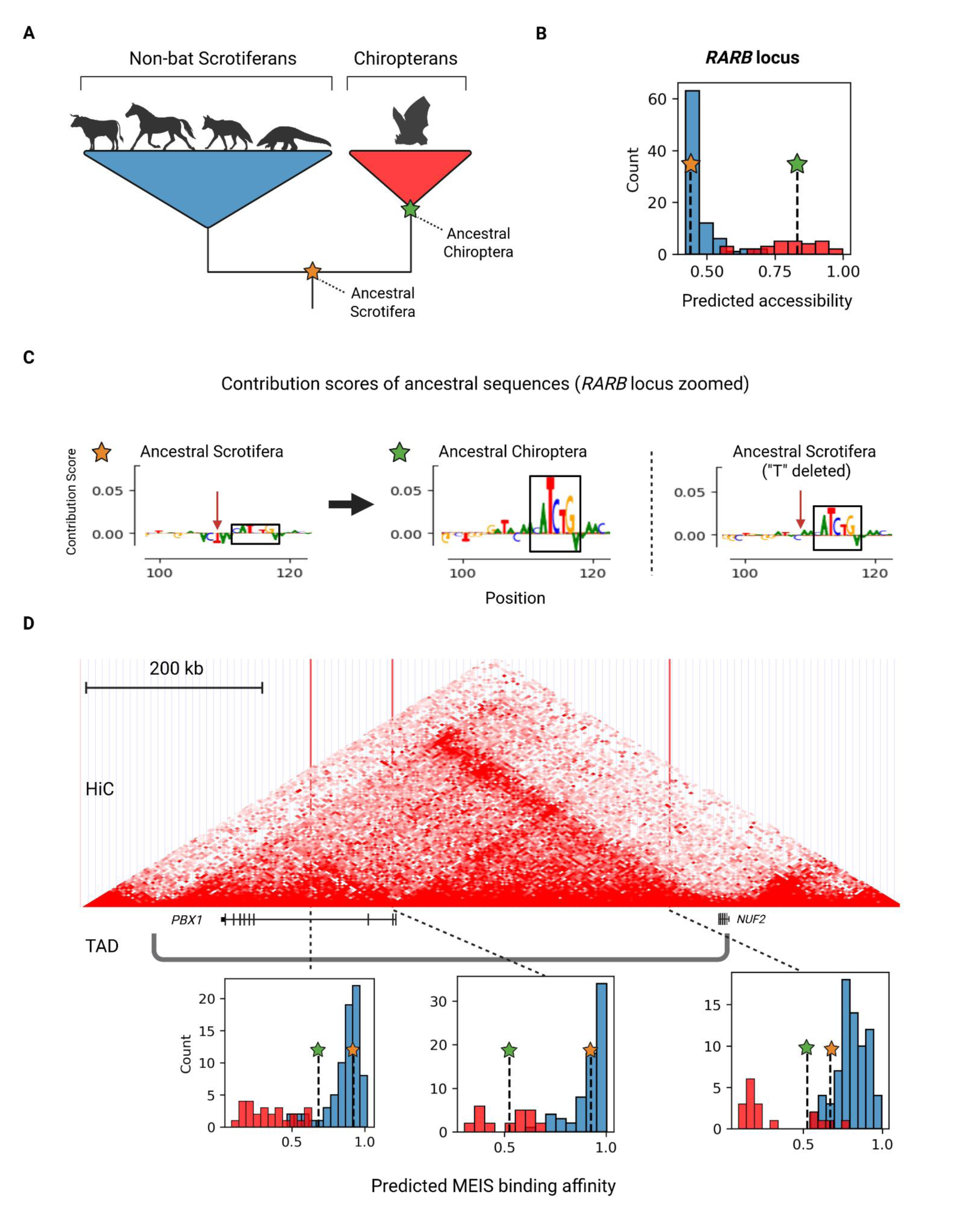
Divergence of regulatory phenotypes near *MEIS2*-interacting genes in bats. **A.** Phylogeny of the clade Scrotifera (superorder: Laurasiatheria). **B.** A *RARB* regulatory element that evolved increased wing-specific accessibility in the ancestral Chiropetra genome, and is maintained in modern bats. Each count represents one species in the Zoonomia genomes **C.** Nucleotide contribution scores of the *RARB* locus. Here, the two ancestral and one *in silico* mutated sequences are shown. The bHLH motif (black squares) in ancestral Scrotifera has lower contribution scores compared to the same motif from ancestral Chiroptera. The difference in scores is partially explained by a deletion (red vertical arrow) in the motif’s flanking region. *In silico* deletion of the nucleotide resulted in a 35% increase in predicted wing-specific accessibility. **D.** Evolution for reduced MEIS affinity in three conserved elements in the *PBX1* TAD. *PBX1* is the only protein-coding gene within this TAD. All three loci (red vertical lines) show contact with the *PBX1* promoter in 3D space in a modern bat genome.

We first reexamined divergence at *cis*-regulatory loci associated with genes upregulated in the bat forelimb (Fig. 5). Previously, we identified forelimb differentially active, *MEIS2*-bound DNA regions in the regulatory domains of these genes. However, whether such distinct activity changes are shared with other bats is unknown. Using the accessibility model to predict wing-specific accessibility (Fig. 3C), we predicted regulatory loci exhibiting bat-specific accessibility divergence patterns. Phylogenetic regression identified 17 loci with divergence in bats (p < 0.05; all above 0.05 after FDR correction), associated with nine putative target genes: *APCDD1*, *BNC2*, *LTBP1*, *OLFML3*, *RARB*, *RARRES2*, *RSPO3*, and *TWIST1* (Supplementary Fig. 3).

*RARB* (retinoic acid receptor beta) was linked to the largest number of divergent loci (five). The gene is crucial for mediating retinoic acid (RA) signaling during limb development (Ghyselinck et al. 1997). RA signaling in the distal limb is an essential component of limb mesenchyme patterning through the induction of interdigital cell death (Dupé et al. 1999). We examined one of these loci with the strongest divergence signal to further explore the details of functional evolution (FDR = 0.1, Fig. 7B). Ancestral sequence reconstruction followed by contribution score analysis revealed increased predicted accessibility in Chiropterans driven by enhanced contributions from a bHLH motif (Fig. 7C). The motif itself is conserved, but a single nucleotide deletion in the flanking region accounts for ∼35% of the accessibility difference between ancestral states, as shown by *in silico* mutagenesis (Fig. 7C). This example illustrates how minimal sequence changes can produce substantial regulatory divergence.

### The flexible evolvability of wing-specific accessibility

Sequence conservation is often interpreted as evidence of functional conservation. However, our analyses show that wing-specific chromatin accessibility can change even within genomic regions where orthologies are traceable across mammalian orders. We therefore examined the evolvability of wing-specific accessibility in conserved sequences. To assess the evolvability of wing-specific accessibility, we applied an algorithm to iteratively mutate unacc. peak orthologs (Fig. 3A). At each iteration, a single nucleotide substitution that maximized the predicted accessibility increase was introduced (Supplementary Fig. 4A). Remarkably, most sequences reached near-maximal accessibility after only five substitutions, and all converged to full accessibility after nine substitutions (Supplementary Fig. 4B). These results indicate that fewer than ten nucleotide changes are sufficient to confer wing-specific accessibility on preexisting elements with MEIS binding potential, highlighting the high evolvability of tissue-specific regulatory programs even within highly conserved genomic regions.

### Evidence of Chiroptera-specific adaptations in the regulatory domain of the MEIS co-factor *PBX1*

Lastly, we assessed genome-wide evolution of *MEIS2* DNA-binding affinity in bats. Because elevated *MEIS2* expression in the bat forelimb is expected to exert selective pressure beyond experimentally detected ChIP-seq peaks, we predicted MEIS binding across 841,603 mammalian conserved elements. This analysis identified 179 elements with significant divergence in predicted MEIS binding (FDR < 0.05, Supplementary Table 6)). To assess the functional relevance of these divergent elements, we performed GO enrichment analysis on their putative target genes assigned by TADs. Enriched terms included system development, biological regulation, and protein binding (Supplementary Table 7). Among the predicted target genes, *PBX1*—a key transcription factor that forms heterodimers with MEIS—was associated with the largest number of bat-divergent elements (three). Hi-C data confirmed contacts between these elements and the *PBX1* promoter in *C. perspicillata* (Fig. 7D). All three loci exhibit reduced predicted MEIS binding affinity in bats, with ancestral reconstructions placing the origin of these reductions coinciding the emergence of Chiroptera. Two of the three loci overlap with regions previously reported to show accelerated evolution in bats (Supplementary Table 6) (Ferris et al. 2018). Together, we reveal coordinated, bat-specific regulatory evolution affecting the MEIS core partner *PBX1*. These results highlight how sequence-based molecular phenotyping enables detection of adaptive regulatory changes associated with morphological innovation.

## Discussion

MEIS genes are key regulators of multiple developmental processes (Schulte and Geerts 2019), including early limb bud formation. In contrast, the later expression of *MEIS2* in the interdigital space has received relatively little attention. Previous studies suggest that interdigital *MEIS2* maintains cell proliferation in mice and contributes to forelimb specialization in bats (Mason et al. 2015; Schindler et al. 2025). Here, we study in detail the role of *MEIS2* in bat forelimb development from both proximate mechanistic and ultimate evolutionary perspectives.

Through cross-species functional genomics integrated with machine learning, we identified frequent and spatially adjacent binding of *MEIS2* with bHLH transcription factors, most likely *TWIST1*, in the bat forelimb. This co-occurrence is further supported by single-cell RNA-seq data, which reveal bat-specific upregulation of *TWIST1* in *MEIS2*-expressing forelimb cells. In humans, *TWIST1* is predominantly expressed in distal mesenchymal tissues and, to a lesser extent, in interdigital regions during digit development (Zhang et al. 2024). Mutations in *TWIST1* cause Saethre-Chotzen syndrome (Ghouzzi et al. 1997; Howard et al. 1997), in which syndactyly is common. In bats, *TWIST1* is upregulated in elongated forelimb digits, although some of these signals may derive from adjacent wing membrane tissues due to the limitation of bulk transcriptomics (Wang et al. 2010). Together, these observations support a role for *TWIST1* in digit and interdigital development, making it a plausible participant in *MEIS2*-mediated wing morphogenesis.

*TWIST1* can mediate tissue-specific regulatory functions through cooperative interaction with other transcription factors. Recent studies in craniofacial and limb development identified a homeodomain–*TWIST1* composite motif associated with the emergence of novel enhancers during evolution (Kim et al. 2024). We observed enrichment of this motif in *MEIS2*-binding regions in the bat forelimb. However, both MEIS and its dimerization partner PBXs belong to the TALE homeodomain family, whose binding motifs diverged significantly from the canonical homeodomain motifs (Penkov et al. 2013). Moreover, aside from PBXs, our RIME proteomic analysis did not detect other transcription factors that co-bind with MEIS. It is therefore unlikely that *TWIST1* directly interacts with *MEIS2*. The precise molecular interplay between *TWIST1* and *MEIS2*, as well as the downstream regulatory consequences of their co-occupancy, remains to be determined.

Beyond *TWIST1*, dozens of genes are upregulated in *MEIS2*-expressing bat forelimb cells. Gene Ontology analysis shows enrichment for functions related to gene regulation, limb development, and extracellular matrix organization. Importantly, the regulatory domains of these genes—including *TWIST1*—exhibit strong *MEIS2*-binding signals (Schindler et al. 2025). These findings support a model in which *MEIS2* functions as a central regulator, activating downstream transcription factors and effector genes during wing membrane development.

A major challenge in studying the genetic basis of bat wing evolution is that this trait is ancient and evolved rapidly over a short evolutionary interval (Upham et al. 2019). Moreover, most comparative functional studies include only one representative bat and one contrast species (Ray and Capecchi 2008; Eckalbar et al. 2016; Schindler et al. 2025), making it difficult to infer the directionality of genomic changes. Many observed differences likely postdate the origin of bat flight. We addressed this limitation by integrating deep learning–based molecular phenotyping across dozens of species and analyzing predicted phenotypes using phylogenetic regression. This approach enabled characterization of the genome-wide *cis*-evolutionary landscape of MEIS DNA-binding in bats. Gene Ontology enrichment of bat-divergent *cis*-regulatory loci mirrors transcriptomic findings, supporting the proposed role of *MEIS2* in wing development. Additionally, evidence of divergence for reduced MEIS binding accompanied by accelerated evolution in *PBX1* regulatory elements were detected in bats. Because MEIS mainly function as activators (Schulte and Geerts 2019), we hypothesize that reduced MEIS-binding affinities at these enhancer loci compensate for increased *MEIS2* dosage in the bat forelimb, representing a derived feedback adjustment within the bat *MEIS2* regulatory network.

Tissue-restricted expression modulation through *cis*-regulatory tuning is thought to minimize deleterious pleiotropic effects in evolution. We explored the substitutional landscape underlying wing-specific chromatin accessibility and found that relatively few nucleotide changes are sufficient to confer wing-specific accessibility on preexisting MEIS binding elements. These findings illustrate how conserved sequences can still acquire lineage-specific regulatory functions, consistent with our phylogenetic regression results.

We also observed striking differences in transposable element (TE) content between conserved and non-conserved *MEIS2* binding peaks, likely reflecting strong purifying selection against TE insertions in conserved regions. In non-conserved regions, however, TEs contribute substantially to MEIS binding. This contribution is not restricted to a particular TE class, suggesting that TEs broadly act as sources of regulatory innovation or mutational hotspots, facilitating the emergence of lineage-specific regulatory elements involved in bat wing development.

Machine learning has rapidly become a powerful framework for studying regulatory genomics (Avsec et al. 2021; de Almeida et al. 2022; Vaishnav et al. 2022). Here, we demonstrate that compact deep learning models trained to predict specific epigenomic features can effectively address evo-devo questions. A limitation of our regulatory element–centric approach is that it can only be implemented to study evolution within conserved regions and does not account for changes occurring elsewhere in the genome. Recent transformer-based deep learning models, which can predict gene expression from sequence contexts spanning megabases (Avsec et al. 2026), offer a complementary, gene-centric perspective. Future studies incorporating additional tissues and developmental stages will provide a more complete view of the molecular mechanisms and the evolutionary forces shaping bat wing development.

In conclusion, by integrating comparative functional genomics, single-cell transcriptomics, and machine learning within a phylogenetic framework, we provide insight into how morphological innovations are regulated during evolution. Together, these findings suggest that the evolution of the bat wing involved coordinated modulation of both transcription factors and genome-wide *cis*-regulatory binding landscapes, rather than isolated changes at individual loci. Our results deepen our understanding of the complex operation and evolutionary trajectory of *MEIS2* during the origin of powered flight.

## Materials and Methods

### MEIS ChIP-seq and ATAC-seq Data Processing

MEIS ChIP-seq data from the autopod region of embryonic forelimb (stage late CS18 for Seba’s short-tailed bat and E15.5 for mouse) were generated in a previous project (Schindler et al. 2025). Additionally, MEIS ChIP-seq data from E10.5 forelimb (SRR9659693-SRR9659696) (Delgado et al. 2021) and adult heart (SRX7299483-SRX7299488) (Muñoz-Martín et al. 2025) were downloaded. The reads were aligned to their respective genomes (carPer2 for bat (Schindler et al. 2025) and mm39 for mouse) using STAR v2.6.1d (Dobin et al. 2013). Alignments were sorted with Samtools v1.22 (Li et al. 2009). Bigwig files normalized by counts per million (CPM) were generated with deepTools v3.5.5 bamCoverage (Ramírez et al. 2014). Reproducible peaks for ChIP-seq, including input controls, were called using Genrich with the parameter -a 50 (Gaspar 2025). In addition, a less stringent set of peaks were called using macs2 (Zhang et al. 2008) to generate negative sets for the MEIS accessibility model and for wing-specific accessibility evolution analysis. ATAC-seq data from the whole forelimb of bat (CS17) and mouse (E10.5), previously published (Paliou et al. 2019; Schindler et al. 2025), were processed using the same pipeline. Peak calling for ATAC-seq was also performed with Genrich using the parameter -j. ChIP-seq data was visualized with IGV (Thorvaldsdóttir et al. 2013).

### Orthology and Conserved Genomic Regions Across Species

Orthologous MEIS regions between the two species were identified using IPP (Phan et al. 2025). Pairwise sequence alignments with ten chordate genomes served as bridging species for IPP. MEIS ChIP-seq summits from E10.5 mouse and CS18 bat were projected onto each other’s genomes. Peaks were labeled as within conserved regions if classified as DC (directly conserved) by IPP, and as non-conserved if labeled IC (indirectly conserved) or NC (non-conserved). Visualization of the local conservation scores of different classes were done by first projecting the bat peak summits onto the mm39 genome with IPP. Deeptools was then used to plot the 35-way vertebrate phastCons and phyloP scores (Casper et al. 2025) of these projected summits and the flanking +-250bp region into heatmaps.

### Motif Enrichment Analysis

*De novo* motif enrichment analysis was performed with Homer2 (Heinz et al. 2010) on the early proximal mouse and late interdigital bat MEIS peaks. Default Homer2 parameters were used, which search the center 400 base pairs of an input region. Random genomic sequences served as background. We also conducted motif enrichment analysis using the built-in homer database. This includes the TWIST1-homeodomain cooperative motif, which was labeled as Pitx1:Ebox(Homeobox,bHLH)/Hindlimb-Pitx1-ChIP-Seq(GSE41591)/Homer in the Homer database.

### Deep Learning Data Preparation: Bat MEIS Binding Model

Three binary classification convolutional neural network models were built for the study. For the bat MEIS binding model, positive labeled data comprised ±200 base pair regions around MEIS peak summits, while negative labeled data included the center 401 base pairs of ATAC-seq peaks not intersecting MEIS peaks. This input size was used to match the motif enrichment analysis window size. Differences in centering for ChIP-seq and ATAC-seq peaks were due to lower coverage at actual transcription factor binding sites in ATAC-seq data. Data augmentation into twofold was done by adding reverse complement sequences generated with SeqKit v2.9.0 (Shen et al. 2024). Data was partitioned into training, validation, and testing sets based on chromosome origin: sequences from chr8 and 10 of the carPer2 genome comprised the validation set; sequences from chr1 were used for testing; the remaining sequences formed the training set. DNA sequences were one-hot encoded into 2D matrices (length × 4), with bases ordered A, C, G, and T.

### Deep Learning Data Preparation: Mouse *TWIST1* Binding Model

*TWIST1* ChIP-seq data came from GSM7213714 (Kim et al. 2024). The Mouse *TWIST1* binding model follows the same design logic as the Bat MEIS binding model, albeit using ATAC-seq and ChIP-seq from E10.5 mouse forelimb. Sequences from chr8 and chr9 of the mm39 genome formed the validation set; chr1 and chr2 formed the test set; all other regions were in the training set.

### Deep Learning Data Preparation: Accessibility Model

For the accessibility model, positive labeled data included bat-specific MEIS peaks, and negative labeled data comprised unacc. peak orthologs. Input sequences were 501 base pairs surrounding either MEIS ChIP-seq summits (positive set) or projected summits from the mouse (negative set). Due to the smaller number of available sequences, data was augmented sixfold with neighboring regions overlapping by 401 base pairs and reverse complements. Sequences projected to chr8 and chr9 of the mm39 genome formed the validation set; chr1 and chr2 formed the test set; all other regions were in the training set.

### Model Building, Training, and Interpretation

Convolutional neural networks were constructed using tensorFlow v2.11.0 keras, employing different model architectures for the three models. Generally, each model comprised two to four convolution layers with ReLU activation, followed by batch normalization, max pooling (size 35). Afterward is a flattening layer, and one fully connected layer (ReLU activation, 0.3 dropout). The final layer was a single node with sigmoid activation. We used the Adam optimizer, binary cross entropy as the loss function, and ROC AUC for evaluation. Early stopping with a patience of three restored the best weights.

Interpretation of models utilized nucleotide contribution scores with DeepExplainer from SHAP v0.41.0. Background sequences for DeepExplainer comprised 100-times dinucleotide-shuffled sequences. Hypothetical contribution scores were translated into actual scores by multiplying the scores with the one-hot encoded sequence and visualized with viz_sequence from DeepLIFT v0.6.12.0. TF-modisco v0.5.16.4.1 was used to identify motifs using actual contribution scores, with parameters set at target_seqlet_fdr=0.2, and max_seqlets_per_metacluster=50000. Predicted motifs were compared to known motifs from the JASPAR database (Ovek Baydar et al. 2025) with tomtom v5.1.1 (Gupta et al. 2007).

### Prediction in different datasets

The bat MEIS binding model was applied to predict MEIS binding within E10.5 mouse accessible genomic regions. The same data processing procedure was used to label the E10.5 mouse peaks. Performance of cross-species predictions was evaluated using ROC AUC. The mouse *TWIST1* binding model was applied to the mouse MEIS ChIP-seq peaks from E10.5 forelimb, E15.5 fore- and hindlimb, adult heart, and bat CS18 forelimb. Due to the high false positive of E15.5 samples, we first filter the E15.5 peaks by the bat MEIS binding model. Only peaks with a MEIS prediction value over 0.7 were used for *TWIST1* prediction.

### Single-cell RNA-seq analysis

Using our previously generated mouse and bat developing limb atlas (Schindler et al. 2025), we extracted cells of interdigital origin. First, we isolated all cells expressing (>0 UMIs) at least one of three well-known autopod markers *HOXD13, HOXA13, MSX1.* These cells were integrated into autopod datasets, and analyzed as previously described. We then further selected cells expressing *MEIS2* which formed part of fibroblastic clusters (clusters expressing *COL1A1*). We compared the gene expression of the datasets’ highly variable genes (normalized variability > median calculated using Seurat function VariableFeatures) in these cells between fore- and hindlimb in both mouse and bat datasets using the MAST implementation in the Seurat v3 function FindMarkers (Hao et al. 2024). The significantly differentially expressed genes (log2FC > 0.25, adj. p.val < 0.1) of both species were then re-tested to avoid specific differences arising from thresholds. Gene ontology enrichment analysis was done with the g:profiler server using the human genome.

### Bulk histone ChIP-seq analysis

Bulk H3K27act ChIP-seq from the autopod region of embryonic forelimb (stage CS19 for Seba’s short-tailed bat were generated in a previous project (Schindler et al. 2025). Forelimb-specific differential H3K27act regions were called with macs2 bdgdiff against hindlimb ChIP-seq using the parameter -l 800 -g 500.

### Use (RIME) for MEIS2 protein interaction in the bat forelimb

Bat samples (*Carollia perspicillata*) were obtained from a captive population maintained at the Papiliorama zoo in Kerzers, as previously reported (Schindler et al. 2025). Bat embryonic forelimbs (staged late CS18) were dissected and fixed in 1% formaldehyde in 10% FCS/PBS for 10 minutes. After quenching in 125 mM glycine, they were snap-frozen and stored at −80 °C. The RIME protocol was adapted for further processing (Mohammed et al. 2016). Briefly, nuclei were extracted, washed, and lysed using LB1, LB2, and LB3 solutions, respectively, before sonication on a Diagenode Bioruptor Plus Sonication device (40 cycles, 30 s on, 30 s off, at high power setting). The lysate was incubated overnight at 4 °C with rotation, with magnetic beads (100 μl Dynabeads Protein A (Invitrogen, #10001D)) and two anti-MEIS antibodies simultaneously. One recognizes the conserved C-terminal domain of MEIS1a and MEIS2a, and the other recognizes all MEIS2 isoforms as previously described (Delgado et al. 2021). Antibodies were generously provided by M. Torres. A total of 5 µg of each antibody was used per immunoprecipitation. IgG-only (10 μg) and bead-only controls were included. The distal part of the forelimb was used as a single replicate, while two proximal parts coming from the same embryo were used for the negative controls. Further processing followed the RIME original protocol until processing for LC-MS/MS analyses. In brief, reduction and alkylation of the cysteines were performed sequentially by adding tris(2-carboxyethyl)phosphine to a final concentration of 5.5 mM for 30 minutes at room temperature, followed by the addition of chloroacetamide to a final concentration of 24 mM for a further 30 minutes on the rocking platform. The proteins were then digested with 100 ng trypsin (Roche, MS grade) and incubated overnight at 37°C in a thermal shaker at 900 rpm and pH 8.0. The peptides were acidified with formic acid to a final concentration of 2%. Five, 10, and 50% of the digests were loaded onto Evotip Pure tips (Evosep, Odense, Denmark) according to the manufacturer’s protocol. Peptide separation was performed using nanoflow reverse-phase liquid chromatography (Evosep One, Evosep) and the Aurora Elite column (15 cm x 75 µm ID, C18, 1.7 µm beads, IonOpticks, Victoria, Australia), employing the Whisper zoom 20-SPD method. The LC system was coupled online to a timsTOF Ultra 2 mass spectrometer (Bruker Daltonics, Bremen, Germany) using data-dependent acquisition (DDA) with parallel accumulation serial fragmentation (PASEF). The MS data were processed using MaxQuant (v2.7.0.0) and searched against the Mus_musculus UniProtKB proteome (UP000000589) as of 11/07/2025 and the carPer2 annotation. The “match between run” feature and label-free quantification were used. The mass spectrometry data have been deposited to the ProteomeXchange Consortium via the PRIDE partner repository. Only proteins with a sample/control ratio of over 100 were considered positive. Two collagen genes were also removed as they are abundant extracellular proteins in the tissue. R package wordcloud2 was used to visualize the results, using the log sample/control ratios as font sizes.

### TE annotation and analysis

A manually curated bat repeat library (Jebb et al. 2020) was used to annotate repeats in the carPer2 genome using RepeatMasker v4.1.7-p1 (Tarailo-Graovac and Chen 2009). Five TE classes (DNA, LINE, LTR, RC, Retroposon, and SINE) were extracted from the repeat annotations. Density of TEs within conserved and non-conserved MEIS binding peaks were visualized with deepTools. The TEs that intersect with the core 401 bp of non-conserved MEIS peaks were randomly shuffled using a custom python script. The TE shuffled peaks were evaluated with the MEIS binding model. Effects of shuffling were calculated with log odds ratio changes (Zhou and Troyanskaya 2015).

### Analysis of wing-specific accessibility evolution in MEIS binding peaks

The carPer2 genome was attached to the Zoonomia genome alignment using cactus v2.5.1 (Armstrong et al. 2020) commands recommended by the cactus manual, replacing the original carPer1 genome. The MEIS binding regions in narrowPeak format were projected to all the Scrotiferan genomes (30 bats, 86 non-bat Scrotiferans) using HALPER -max_frac 2 -min_len 50-protect_dist 5 (Zhang et al. 2020). Wing-specific accessibilities were predicted on the central 501 base pair region of each peak using the accessibility model. Mean scores were taken from predictions in both strands. Predictions were also made on two reconstructed inner nodes in the alignment: ancestral Chiropteran (fullTreeAnc150) and ancestral Scrotiferan (fullTreeAnc236). TADs for carPer2 were obtained from our previous publication based on Hi-C sequencing from fibroblasts (Schindler et al. 2025). Peaks that are located within TADs that contain the 39 genes found in our single-cell transcriptome analysis, and had an increased prediction in ancestral bat relative to ancestral Scrotifera were identified. The segregation of accessibilities in these peaks between bats and non-bat Scrotiferans was tested using the phylogenetic regression Pagel’s lambda model (Pagel 1999) implemented in phylolm v2.6.5 (Tung Ho and Ané 2014). The input tree was obtained from the Zoonomia website. FDR was used to correct for multi-testing. Nucleotide contribution scores were estimated for the two ancestral sequences. A third, single-nucleotide-deleted ancestral Scrotifera sequence was created manually. Prediction with accessibility model and interpretation by contribution scores were done on this sequence.

### Analysis of MEIS Binding Evolution in Mammalian Conserved Elements

Coordinates for 1,612,714 conserved elements were obtained from a published study on 120 mammalian genomes (Hecker and Hiller 2020). Elements shorter than 100 base pairs were excluded, resulting in 841,603 elements. These filtered elements were projected from the hg38 genome to the Scrotifera genomes within the Zoonomia 241 genome alignment using HALPER. Only 170,158 elements, which were projected to at least 15 bat genomes, were kept. MEIS binding predictions were made on the central 401 base pair region of each conserved element using the bat MEIS binding model, with mean prediction scores taken from both strands. The segregation of molecular phenotypes between bats and non-bat Scrotiferans was assessed using phylolm. FDR was applied to correct for multiple testing. The number of bat-divergent elements within each TAD was calculated using Bedtools v2.30.0 intersect (Quinlan and Hall 2010). The Hi-C contact matrix was visualized with the UCSC genome browser. Genes that are located in the TADs containing the bat-divergent elements were used for GO enrichment analysis with the g:profiler server (Raudvere et al. 2019) using the human genome as a reference. The keratin genes were excluded from the analysis as over twenty of them occur in tandem arrays inside a TAD that contains one bat-divergent element. The three *PBX1*-related elements were projected to the hg19 genome and then intersected with published bat accelerated regions (Ferris et al. 2018).

### Genetic algorithm and simulating evolution

A genetic algorithm that exhaustively searches and returns the single nucleotide substitution with the largest gain in prediction score was written in python. All non-MEIS peak orthologs of mouse peaks in the carPer2 genome were fed to the algorithm. The genetic algorithm was applied for 10 iterations to simulate selection favoring increased wing-specific accessibility. The sequences whose accessibility cannot be increased by any possible single nucleotide substitutions in the first iteration were excluded from the analysis.

## Data Availability

Supplementary information can be found online. All sequencing data came from published datasets (see Materials and Methods). The RIME results are deposited in the PRIDE database under the accession PXD074776. The deep learning models and instructions for using them are available in: https://github.com/BaiweiLo/Bat-MEIS-models.

## Authors’ contributions

**B. L.** designed the study, conducted the machine learning and genomic analyses. **S. A.** collected and prepared the biological samples for RIME. **D. M.** perform the protein mass spectrometry. **C. F.** performed scRNA-seq analyses. **M. S.**, **F. M. R.**, and **A. R.**, assisted with sample collection. **B. L.** drafted the manuscript. **B. L.** and **C. F.** finalized the manuscript. **S. M.** provided funding. **C. F.** and **S. M.** supervised the study.

## Acknowledgements

We thank Natalia Benetti for her input on the manuscript. This work has received funding to S. M. from the European Research Council (ERC grant GenRevo, No. 101054341).

